# Epitranscriptome analysis of HeLa-expressed HPV18 early transcripts

**DOI:** 10.1101/2025.07.28.667157

**Authors:** Ida Rye Hellebek, Malte Storm Lau Schlosser, Johanna Jönsson, Jamie Yam Auxillos, Sarah Rennie

## Abstract

Human papillomavirus (HPV)-driven cancers remain a major global health burden, and understanding how post-transcriptional regulation shapes viral gene expression may inform new therapeutic strategies. Here, we reanalysed independently generated public HeLa datasets to identify candidate RNA modification sites on integrated HPV18 transcripts expressed from the HeLa genome. Using Oxford Nanopore direct RNA sequencing of native RNAs and in vitro transcribed controls, together with GLORI, eTAM-seq and staged 4sU-GLORI datasets, we identified a set of candidate m^6^A sites on HPV18 early transcripts. m^6^A levels at a subset of these sites were reduced following perturbation of the m^6^A pathway, most clearly after METTL3 inhibition, WTAP knockdown or FTO overexpression. We further show that the E6*I-proximal m^6^A site at position 224 is enriched on unspliced transcripts in direct RNA sequencing, eTAM-seq and staged 4sU-GLORI data. In contrast, we find no convincing evidence for m5C or pseudouridine within the HPV18 regions covered by these datasets. Together, our analyses provide a candidate map of RNA modifications on HeLa-expressed HPV18 early transcripts.

## Introduction

Human papillomavirus (HPV) is a small, non-enveloped DNA virus with a circular genome of approximately 8 kb in length (Zhang et al., 2025). HPV is divided into high- and low-risk types based on their association with cancer (Doorbar et al., 2012). Of the various HPV types, HPV18 and HPV16 are the two most prevalent high-risk types, together accounting for ∼70% of HPV-related cervical cancers worldwide, as well as a rising fraction of other cancers such as oropharyngeal carcinomas (de Martel et al., 2017). Despite the availability of vaccines and cervical screening programmes, HPV-associated cancers remain a global health burden, underscoring the need for continued research (Sung et al., 2021; Schiffman et al., 2016).

HPV-driven oncogenesis depends primarily on the sustained expression of the viral E6 and E7 oncoproteins. E7 promotes cell-cycle re-entry, leading to unscheduled DNA synthesis and cell proliferation, whereas E6 promotes the degradation of the tumour suppressor p53 and targets cellular pathways that favour evasion of apoptosis and immortalisation (Doorbar et al., 2012). Later in the viral infection process, accumulation of the E2 protein leads to a down-regulation of E6 and E7 expression and results in the expression of late genes (Doorbar et al., 2012). However, in many long-term HPV-associated cancers, viral integration disrupts E1/E2, reducing E2-mediated repression and contributing to the continuous expression of E6 and E7 (Nishimura et al., 2000; Molina et al., 2024). For example, modern HeLa cells harbour multiple integrated copies of HPV18, where viral-host chimeric transcripts appear to arise from a single transcriptionally active copy and utilise a host polyadenylation signal (Yu et al., 2024). These viral transcripts often read through the E1 coding region, which encodes the viral replicative helicase, and whose expression in HeLa has been observed to affect the expression of immune response genes (Castillo et al., 2014). The E6/E7 mRNA is bicistronic and contains an intron within the E6 open reading frame. Splicing of this intron generates the E6*I isoform, which is required for efficient translation of E7, whereas retention of the unspliced transcript allows continued expression of full-length E6 protein (Tang et al., 2006).

Gene expression is heavily regulated post-transcriptionally by a diverse set of covalent chemical modifications which affect RNA nucleotides, impacting processing, stability and translation (Delaunay et al., 2024; Roundtree et al., 2017). In messenger RNA, the most prevalent internal modification is N6-methyladenosine (m^6^A), which typically occurs at the consensus motif DRACH (D=A/G/U, R = A/G, H= A/C/U). m^6^A is deposited onto RNA by the “writer” core complex, including METTL3, which catalyses the transfer of the methyl group to the adenine base, and METTL14 which assists in substrate RNA recognition and positioning. This process is assisted by WTAP or other factors, such as VIRMA, RBM15 and ZC3H13 (Ping et al., 2014). These marks are additionally read by reader proteins that have a large influence on RNA fate (Zaccara et al., 2019), the best characterised being the YTH-domain family, with others, such as members of the IGF2BP family, acting as additional readers (Huang et al., 2018). In addition to m^6^A, two further modifications occurring on mRNA are 5-methylcytosine (m^5^C) and pseudouridine (Ψ). These are installed by distinct writer enzymes (e.g. NSUN-family enzymes or DNMT2/TRDMT1 for m^5^C, and PUS-family enzymes for Ψ) and have been widely implicated in regulating RNA processing and translation (Song et al., 2022; Liu et al., 2022; Borchardt et al., 2020).

Viruses are able to exploit the molecular machinery of host cells to regulate viral transcription via RNA modification. For example, m^6^A has been implicated in the replication cycle of human immunodeficiency virus (HIV-1) (Lichinchi et al., 2016). In the context of HPV, it was recently shown that the E7-containing HPV16 oncotranscripts are subject to m^6^A methylation and that this mark can increase E7 mRNA stability by recruiting the m^6^A-binding protein IGF2BP1 (**?**). These findings highlight the importance of RNA modifications in shaping virus–host interactions and raise the question of how epitranscriptomic marks influence the regulation of HPV transcripts and oncogenic potential.

Whilst transcriptome-wide detection of precise modified positions has historically been challenging, the recent development of third-generation sequencing has enabled the direct mapping of RNA modifications by sequencing full-length native RNA molecules. Nanopore-based direct RNA sequencing (DRS) detects changes in ionic current as RNA passes through the pore (Leger et al., 2021; Garalde et al., 2018). As the ionic current signal is altered in the presence of chemical base modifications, specialised bioinformatic tools can be used to map modifications on a per-molecule basis (Pratanwanich et al., 2021; Kovaka et al., 2025; Gamaarachchi et al., 2020; Hendra et al., 2022). In practice, samples incorporating *in vitro* transcribed (IVT) mRNA (which lacks endogenous modifications) or writer knockdown controls are often employed to reduce false positives (Tan et al., 2024a; Acera Mateos et al., 2024; Zou et al., 2025), and many studies additionally rely on a combination of approaches to reliably identify sites (Zhong et al., 2023). Recent high-throughput chemical- and enzyme-assisted assays have also been developed to profile specific modifications. For example, GLORI allows for detection and quantification of m^6^A sites by chemically converting unmodified adenosines to inosine (Liu et al., 2023), eTAM-seq detects and quantifies m^6^A by enzyme-assisted adenosine deamination (Xiao et al., 2023), and for m^5^C, m^5^C-TAC-seq was recently developed (Lu et al., 2024).

Because HeLa cells constitutively express integrated HPV18 early transcripts, public HeLa transcriptome datasets provide a practical resource for mining viral RNA modifications without requiring a dedicated HPV infection model. In this study, we reanalysed multiple publicly available HeLa datasets to identify candidate RNA modification sites on integrated HPV18 transcripts. We first used RNA004 nanopore direct RNA sequencing, together with *in vitro* transcribed controls to identify candidate m^6^A, m^5^C and Ψ sites. We then used GLORI, eTAM-seq and staged 4sU-GLORI datasets to prioritise sites that are detected across orthogonal assays. These analyses support a consensus set of candidate m^6^A sites on HeLa-expressed HPV18 early transcripts. In addition, we identify an association between E6 intron retention and methylation at the E6*I-proximal position 224, with methylation preferentially detected on unspliced molecules and in later-stage RNA fractions. Finally, we find little evidence for m^5^C or Ψ in the covered HPV18 transcript regions under the conditions analysed.

## Results

### Direct RNA sequencing provides high coverage of HeLa-expressed HPV18 early transcripts

To assess RNA modifications on HPV18 transcripts, we re-analysed a recently published direct RNA sequencing (DRS) dataset generated from HeLa cells (Zou et al., 2025). This dataset was selected for two key reasons. First, it was generated using Oxford Nanopore’s newer RNA004 direct RNA sequencing chemistry, which provides improved throughput and read accuracy compared to earlier versions. Second, it includes both native RNA from wild-type (WT) HeLa cells and an *in vitro* transcribed (IVT) control, where the IVT sample serves as an unmodified baseline. Our analysis followed the workflow outlined in Figure 1A, using the Oxford Nanopore Dorado basecaller with modification-aware models to obtain candidate m^6^A, m^5^C, and pseudouridine (Ψ) calls, followed by alignment of the reads to a hybrid reference genome (human GRCh38 plus HPV18).

**Figure 1.**
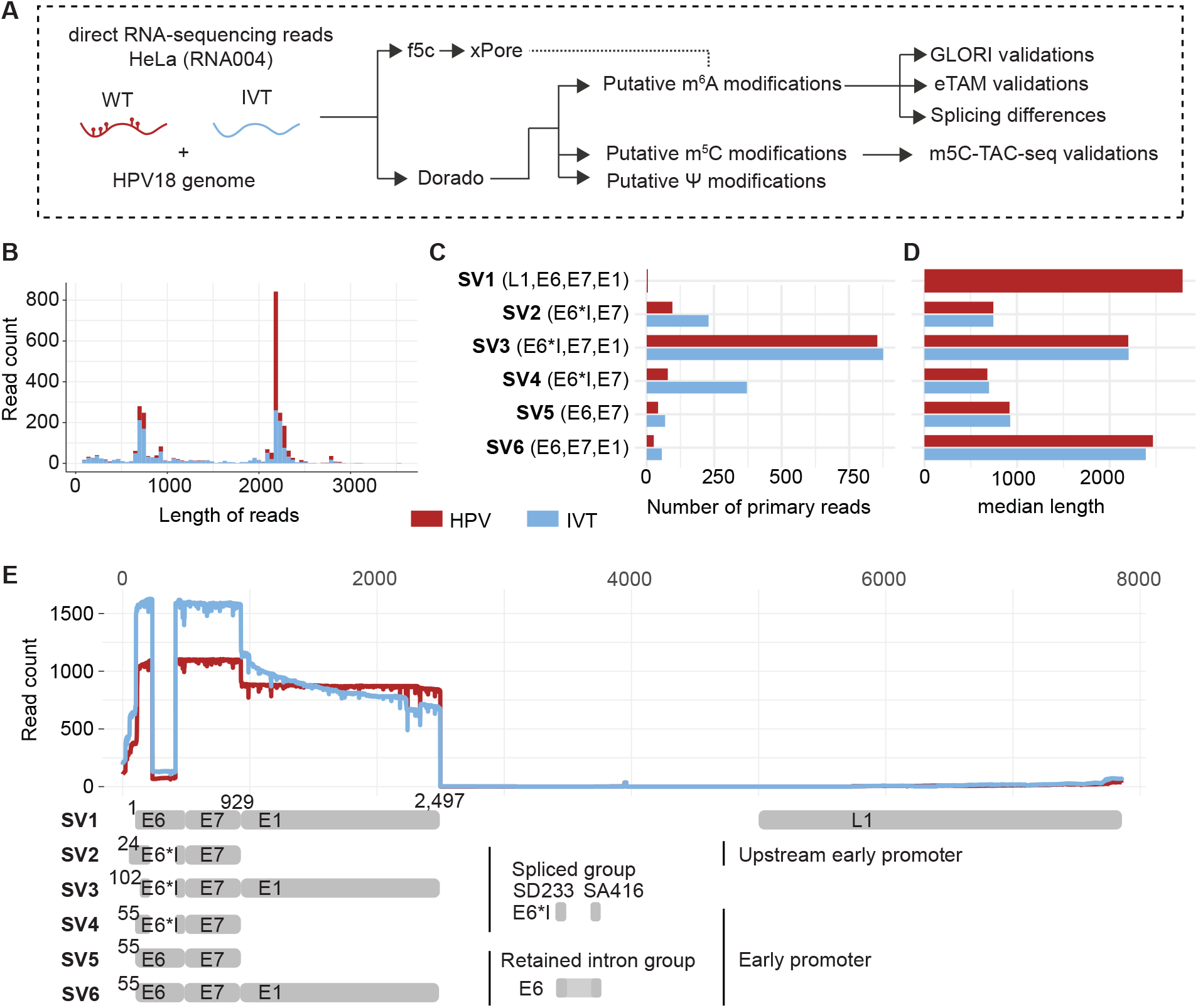
Nanopore direct RNA sequencing reads cover HeLa-expressed HPV18 early transcripts. (**A**) Schematic of the overall study design and analysis workflow. (**B**) Length distribution of DRS reads mapping to the HPV18 genome, shown separately for the WT and IVT samples. (**C**) Number of reads mapping to each of six splice variant (SV) categories. (**D**) Median read lengths for each splice variant category. (**E**) Coverage across the HPV18 genome by nanopore reads, including schematic illustrating the six splice variant categories.

The HPV18-aligned read lengths formed two clusters (approximately 0.8 kb and 2.3 kb) for both IVT and WT samples (Figure 1B). The shorter reads most likely correspond to E6/E7 transcripts, whereas the longer reads correspond to transcripts incorporating the E6, E7, and E1 regions. Almost all HPV18-mapped reads contained soft-clipped host-derived sequence visible in IGV (Figure S1A), and these soft-clipped portions were excluded from downstream analyses. By examining the presence of viral exons and the 5^*′*^ alignment start positions of individual reads, we classified the reads into six splice variant (SV) categories, which were present in both IVT and native WT read sets (Figure 1C). Five correspond to isoforms identified in a previous study (Yu et al., 2024), while a sixth (SV6) was newly defined here, retaining both the E6 intron and the E1 region. The median read length for each splice variant was consistent with expectations (Figure 1D). Furthermore, the coverage profile across the HPV18 genome confirmed high read depth over the E6 and E7 coding regions but low coverage elsewhere (Figure 1E), and the quality of the data was generally good in these high-coverage regions (Figure S1B). These analyses indicate that DRS reads map with sufficient quantity and quality to support downstream analysis of the HeLa-expressed HPV18 early transcript region.

### Dorado identifies multiple candidate m^6^A sites on HPV18 transcripts

We next extracted per-position modification information from the HPV18-aligned reads by aggregating Dorado modification probabilities across reads. Dorado/Modkit reported numerous candidate sites for each of the three queried modification types (m^6^A, m^5^C, and Ψ) in both the WT and IVT samples (Figure 2A). Of the three modification classes, only m^6^A showed substantially more candidate sites in WT than in IVT. By contrast, the numbers of candidate m^5^C and Ψ sites in IVT were similar to, or higher than, those observed in native WT RNA, suggesting that many of these calls may reflect background signal rather than endogenous modifications.

**Figure 2.**
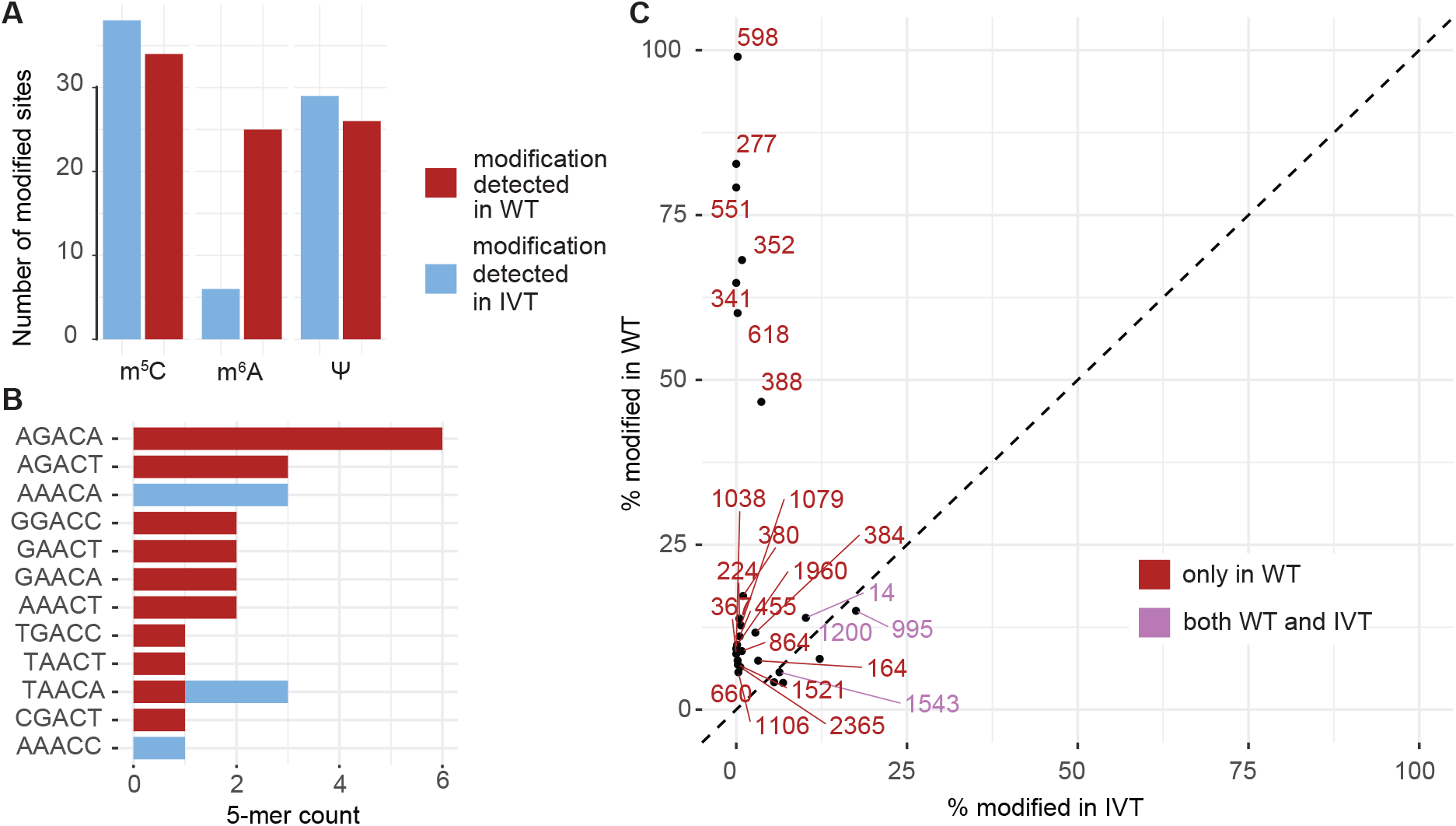
m^6^A modification supported by Dorado-based modification calling. (**A**) Number of candidate sites called in HPV18 for each of m^5^C, m^6^A and pseudouridine, split by WT and IVT samples. (**B**) 5-mers centred on the putative m^6^A sites. (**C**) Scatter plot showing modification rates in WT versus IVT for Dorado/Modkit-identified sites. Red depicts sites with a modification rate greater than 5% in WT; purple depicts sites with a modification rate greater than 5% in both samples.

Focusing on m^6^A, Dorado identified 25 candidate sites in WT HPV18 transcripts, whereas only six sites were called in the IVT sample (Figure 2A), of which four were found in both (Figure S2A). In support of the WT candidates, we observed that the number of these sites detected per individual molecule was much greater in WT than in the IVT control (Figure S2B), and that they typically occurred in common variants of the DRACH motif (Figure 2B). Next, we compared the m^6^A stoichiometries for each site between WT and IVT samples, where stoichiometry is defined here as the proportion of reads classified as modified after applying probability cut-offs in Modkit (Oxford Nanopore Technologies) (Figure 2C). This analysis revealed a subset of sites with substantially higher modification stoichiometry in native WT RNA than in IVT, suggesting that these sites may represent true m^6^A sites in the viral transcripts.

Because Dorado modification calling may depend on model assumptions and sequence context, we used xPore as an independent signal-based analysis to assess support for candidate m^6^A sites (Pratanwanich et al., 2021) (Figure S2C–F). This identified a smaller number of positions that exactly matched the Dorado candidates, but a larger number of WT-specific Dorado m^6^A candidates (15 out of 21) had an xPore-significant position within a ±2 nt window around the Dorado-called site (Figure S2C), consistent with local signal displacement around modified bases. This was significantly more than expected for randomly selected positions tested with xPore (Figure S2D). Consistent with this, manual inspection of event-level signal alignments based on the tool f5c supported two representative positions, 277 and 551 (Figure S2E,F) (Gamaarachchi et al., 2020). Taken together, these analyses support a subset of higher-confidence candidate m^6^A sites on HeLa-expressed HPV18 transcripts for follow-up orthogonal analyses.

### Orthogonal chemical profiling supports a recurrent subset of m^6^A candidates

We next assessed whether the 25 candidate m^6^A sites were supported by independent chemical-profiling data. We first reprocessed published HeLa GLORI datasets (Liu et al., 2023), including the HPV18 reference genome during alignment, and calculated the fraction of protected adenosines at each candidate site. Examination of the GLORI control samples, using the hypoxia^*−*^ control condition because it had the highest read depth, showed substantial coverage across the HPV18 transcriptome (Figure S3A). For each of the 25 Dorado-called candidate sites, we calculated the estimated modified fraction, with many sites showing modification levels consistent with the stoichiometries estimated from direct RNA sequencing data (Figure 3A). For the 21 WT-specific m^6^A candidates, GLORI-derived methylation fractions correlated strongly with the nanopore-based estimates (r^2^ = 0.96, Figure S3C), despite being derived from independent HeLa datasets. In contrast, IVT-supported sites showed protected-adenosine fractions only marginally above the background observed in matched samples not subjected to GLORI chemical conversion (Figure S3B).

**Figure 3.**
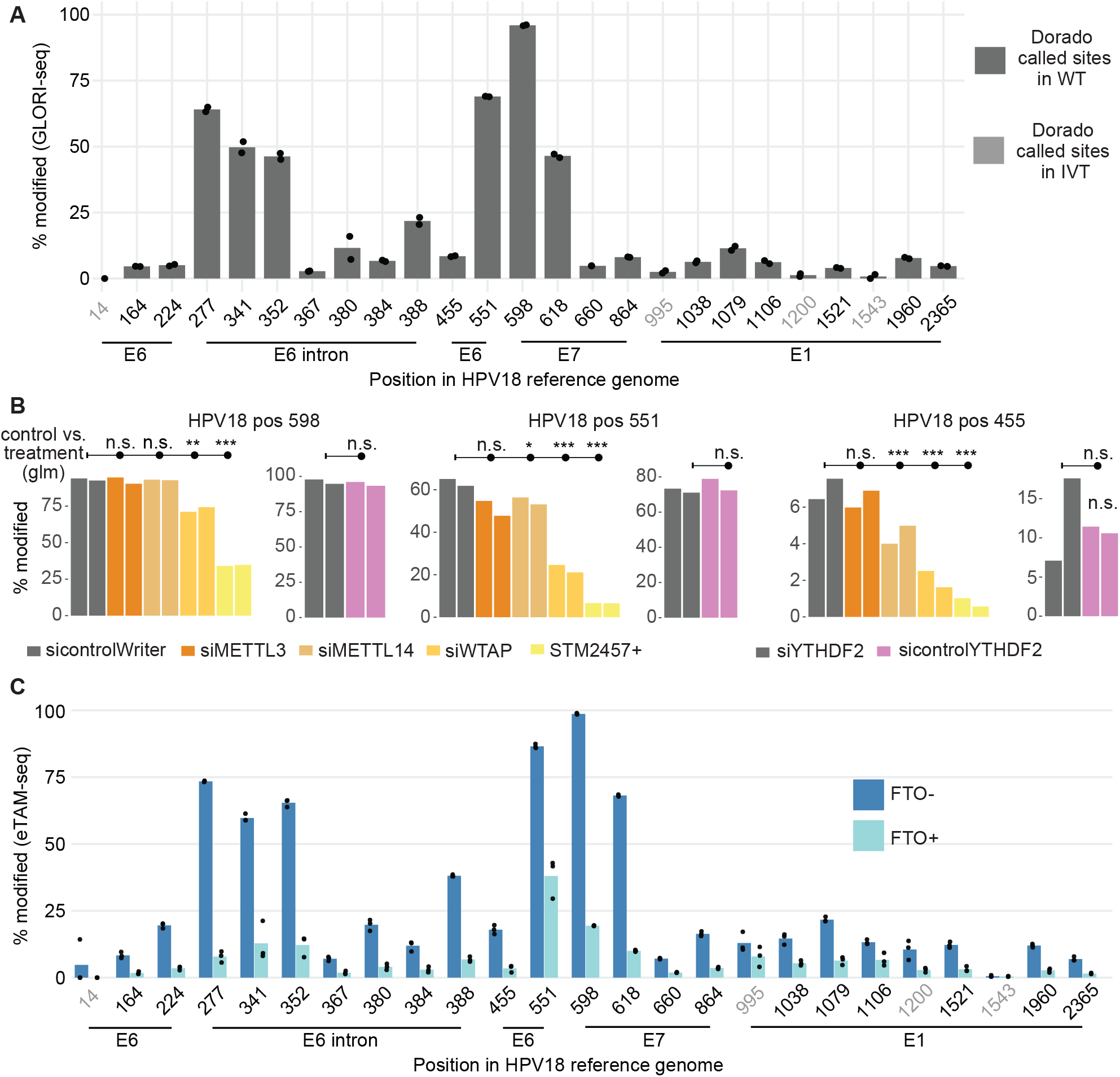
Orthogonal support for Dorado-identified m^6^A sites. (**A**) Fraction of adenosines protected from A-to-I conversion (estimated m^6^A fraction) at each of the 25 Dorado-identified m^6^A candidate positions, as measured in GLORI HeLa datasets. Grey numbers indicate the four candidates that were also detected in the DRS IVT sample. (**B**) Candidate sites 598, 551 and 455 in GLORI following perturbation of the m^6^A writer complex, as well as the reader YTHDF2. Dual bars per position represent the two replicates from the GLORI datasets. Bars above indicate the significance of each perturbation, based on binomial logistic regression models, compared to the grey control samples. Key: n.s., not significant; * p-value < 0.05; ** p-value < 0.01; *** p-value < 0.001. (**C**) Estimated modification percentages from eTAM-seq HeLa data, with or without overexpression of the m^6^A eraser FTO.

Reanalysis of the GLORI perturbation datasets provided additional site-dependent support for involvement of host m^6^A machinery, where the clearest decreases were observed after STM2457 treatment (METTL3 inhibition) and WTAP knockdown (Figure 3B, Figure S4A). For example, position 598 was almost fully methylated in control GLORI samples but decreased after METTL3 inhibition and WTAP depletion. Similar trends were observed at position 551, while position 455 showed lower baseline stoichiometry but was also reduced in some perturbation conditions. These data are suggestive of the involvement of the host m^6^A pathway at candidate HPV18 sites, with the limitation that not all sites had sufficient coverage, and not all writer perturbations produced significant decreases (Figure S4A). In contrast, none of the 25 candidates from direct RNA sequencing showed a significant change in methylation upon YTHDF2 knockdown relative to controls (Figure 3B, Figure S4A), suggesting that loss of this reader protein does not appreciably affect the presence of the methylation marks. Additionally, some sites showed altered m^6^A levels under hypoxic conditions (Figure S4C).

We additionally reanalysed eTAM-seq data from HeLa cells with and without overexpression of FTO, an m^6^A eraser/demethylase. In the FTO^*−*^ control samples, estimated modification fractions at HPV18 candidate sites closely resembled those observed by GLORI (Figure 3C). Upon FTO overexpression, the estimated modification fractions decreased at many sites, consistent with these signals reflecting m^6^A-sensitive adenosines (Figure 3C). The strongest sites, including position 598, showed marked reductions after FTO overexpression, further supporting a consistent subset of HPV18 m^6^A candidates across independent chemical-profiling assays. Taken together, these results indicate support of DRS-detected HPV18 m^6^A candidate sites by GLORI, and further suggest that deposition at at least some of these sites depends on the host m^6^A writer complex.

### Consensus m^6^A candidates across datasets and RNA processing stages

Intersecting the DRS, GLORI and eTAM-seq candidate sets identified 16 consensus m^6^A sites detected by all three approaches (Figure 4A). We next reanalysed staged nuclear 4sU-GLORI data from HeLa cells to ask whether these consensus HPV18 m^6^A candidates varied across nuclear RNA-processing states (Tang et al., 2024). For the 16 consensus sites, modified fractions generally increased from the early 4sU^+^ poly(A)^*−*^ state to the mid 4sU^+^ poly(A)^+^ state and the late 4sU^*−*^ poly(A)^+^ state (Figure 4B). This pattern is consistent with the original 4sU-GLORI study’s observation that many m^6^A sites accumulate post-transcriptionally or become enriched in later nuclear RNA-processing states. Several sites in or near the E6 intron, together with positions 551 and 618, showed particularly clear stage-dependent increases. In contrast, position 598 was already highly methylated by the mid-stage fraction, suggesting that individual HPV18 m^6^A sites may differ in the timing or persistence of methylation. After stringent filtering, we also identified eight additional sites detected by both GLORI and eTAM-seq but not by Dorado (Figure 4A). Although we treat these as lower-confidence candidates, their absence from the DRS candidate set may reflect lower stoichiometry, non-canonical sequence context and/or limited DRS coverage at the relevant positions (Figure 4D). For example, position 932 showed a GLORI modification level of approximately 21% (Figure 4C) but lies outside a canonical DRACH context (GTACA), while several other sites occur in regions with very low DRS read depth. These sites are therefore reported as additional candidates, although subsequent analyses focus on the core set of 16.

**Figure 4.**
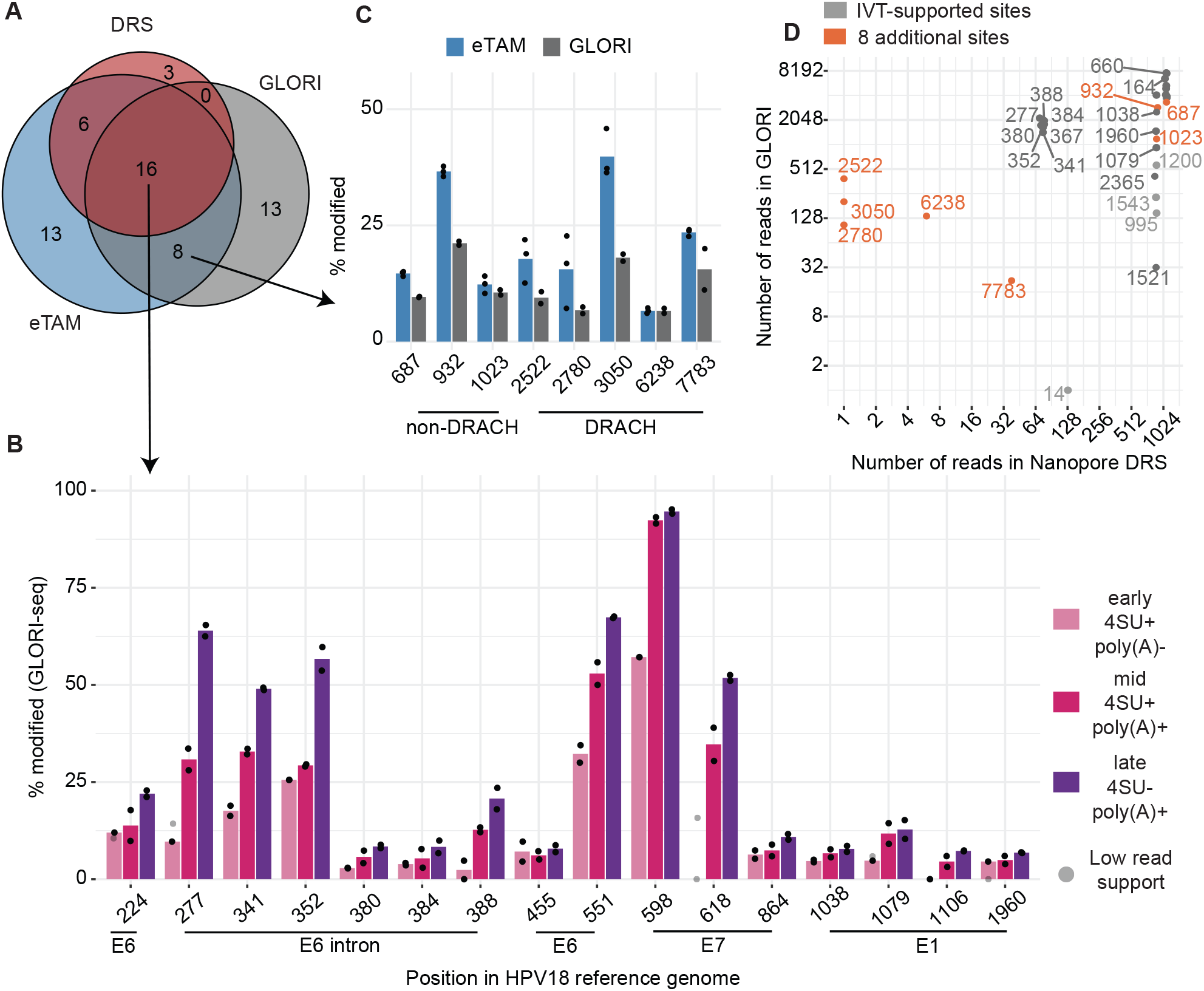
Consensus and additional HPV18 m^6^A candidates across orthogonal assays. (**A**) Venn diagram showing agreement between m^6^A sites identified in HPV18 transcripts in DRS, GLORI and eTAM-seq HeLa data. (**B**) Protected-A proportions in staged 4sU-GLORI data for the 16 sites identified in all three DRS, GLORI and eTAM-seq analyses. Black dots indicate replicate values with sufficient coverage; light grey marks indicate site/stage combinations with insufficient read support. (**C**)Protected-A proportions for eight additional sites detected by both GLORI and eTAM-seq but not by Dorado. (**D**) Read coverage at all DRS-derived candidate positions and the additional eight GLORI/eTAM-supported sites. Colours indicate sites detected in DRS WT only, in both WT and IVT, or in the additional GLORI/eTAM-supported set.

### m^6^A at the E6*I-proximal site is enriched on unspliced HPV18 transcripts

One advantage of long-read direct RNA sequencing is the ability to assess RNA modifications in the context of specific splice isoforms. We therefore asked whether m^6^A stoichiometry differed between HPV18 reads retaining the E6 intron and reads corresponding to the spliced E6*I isoform. DRS reads were divided into E6 intron-retaining reads, which preserve the full-length E6 open reading frame (the unspliced group, 62 reads), and E6*I-spliced reads (the spliced group, 1005 reads) (Figure 5A).

**Figure 5.**
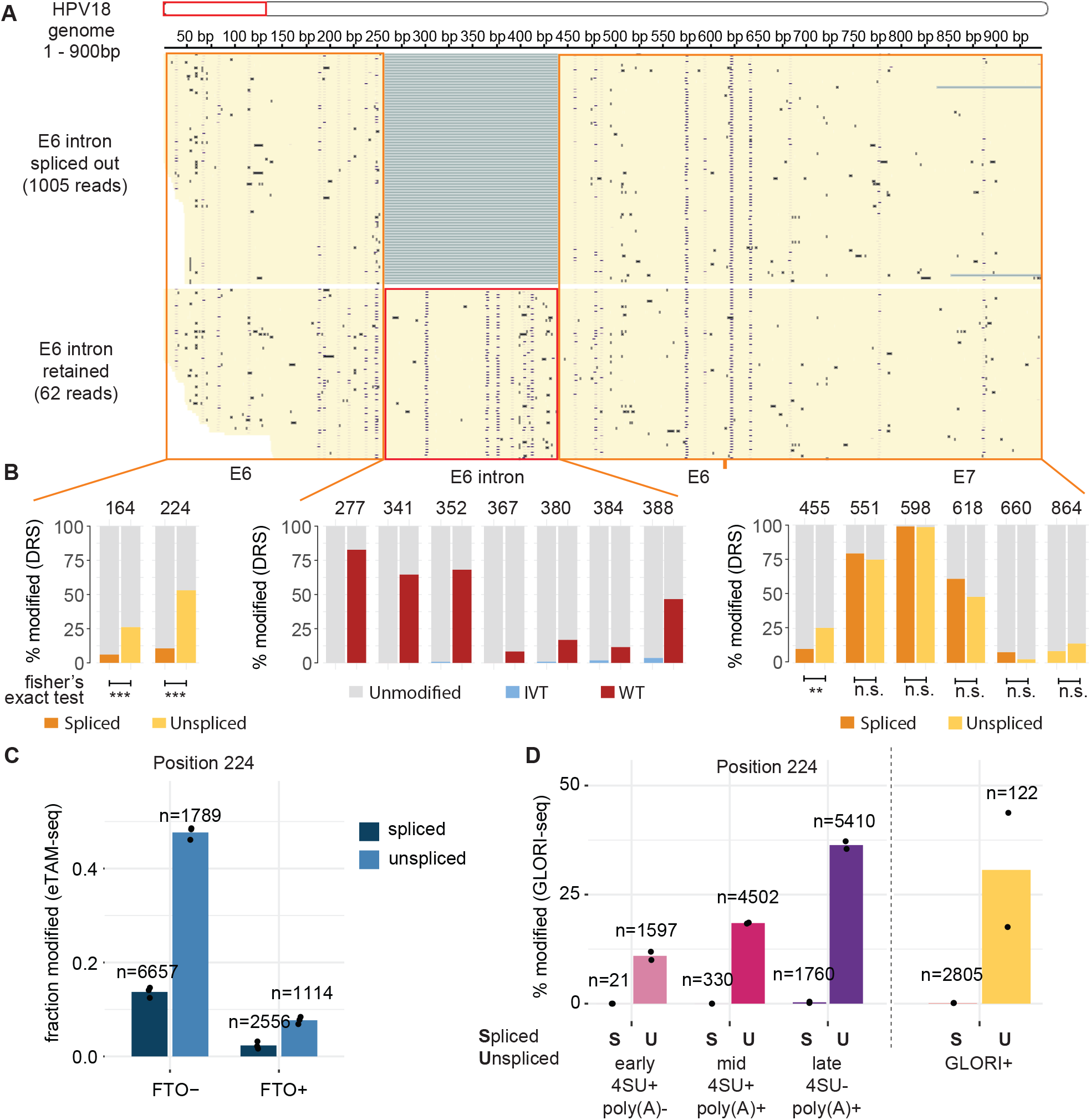
E6 intron splicing status is associated with m^6^A levels at the E6*I-proximal position 224. (**A**) IGV browser view of the first approximately 900 nt of the HPV18 reference, split into reads corresponding to E6*I-spliced transcripts and reads retaining the E6 intron. Blue dots represent Dorado m^6^A modification probabilities. (**B**) DRS modification proportions for high-confidence candidate m^6^A sites in the E6/E7 region, stratified by splice status where informative. Fisher’s exact tests compare modified and unmodified read counts between spliced and unspliced read classes. (**C**) Linkage analysis of eTAM-seq reads covering position 224 and informative for E6 splicing status, shown for FTO- and FTO+ samples. (**D**) Linkage analysis of staged 4sU-GLORI and control GLORI reads covering position 224 and informative for E6 splicing status. n values indicate the number of informative reads contributing to each estimate. Key: n.s., not significant; * adjusted p-value < 0.05; ** adjusted p-value < 0.01; *** adjusted p-value < 0.001.

Within the early region (∼1–900 bp), we observed two m^6^A candidate sites upstream of the E6 splice donor, positions 164 and 224, with higher estimated methylation in unspliced reads than in spliced reads (Figure 5B). This difference was strongest at position 224, which lies in a GAACT motif immediately upstream of the splice donor. Downstream of the E6 intron acceptor, position 455, which lies in an AAACT context 40 bp downstream of the acceptor, also showed evidence of splice-status-associated methylation, although the difference was less pronounced than at position 224 (Figure 5B). Importantly, none of the putative m^6^A sites considered in this analysis showed appreciable modification in the IVT control, even after stratification by splice status, indicating that the observed differences are unlikely to reflect sequencing artefacts (Figure S5B).

To test whether this isoform-linked pattern was also observed in independent short-read chemical-profiling datasets, we performed read-level linkage analyses for reads covering position 224 and informative for E6 splicing status (Figure 5C,D). In eTAM-seq, position 224 showed higher modified fractions in unspliced reads than in spliced reads in the FTO^*−*^ condition, while FTO overexpression showed the same relationship but with reduced modification fractions in both read classes (Figure 5C). In staged 4sU-GLORI data, the modified fraction at position 224 was almost zero in spliced reads at all stages, whereas unspliced reads showed increasing modification fractions across RNA-processing stages (Figure 5D). A similar, but less distinct, pattern was observed for position 455 (Figure S6A,B). We additionally examined E6 intron retention and splicing indices in the eTAM-seq, 4sU-GLORI, and GLORI writer-perturbation datasets (Figure S6C–E). Although these analyses rely on short reads and should therefore be interpreted cautiously, they suggest that E6 intron splicing metrics vary with FTO-dependent eraser perturbation (Figure S6C), GLORI-assayed writer-complex perturbation (Figure S6D), and RNA-processing state (Figure S6E). These observations are broadly consistent with a recent report that perturbation of m^6^A machinery influences HPV16 mRNA splicing (Cui et al., 2022).

Finally, we explored whether other transcript features might also be associated with differential m^6^A deposition. Specifically, we examined (1) transcripts that include the E1 region versus those that do not, and (2) transcripts initiating from an earlier cryptic start site at +24 versus others (SV2 versus others, Figure 1E, (Yu et al., 2024)), to ask whether any m^6^A sites showed analogous isoform-specific patterns. In both cases, we did not detect clear differences in m^6^A levels comparable to those seen for E6 intron retention. Overall, these results identify position 224 as the clearest splice-status-associated m^6^A candidate site, with methylation preferentially enriched on transcripts retaining the E6 intron.

### Little evidence for m^5^C or pseudouridine modifications in HeLa-integrated HPV18 transcripts

As shown earlier, Dorado detected a number of candidate 5-methylcytosine (m^5^C) sites in both WT and IVT samples on HPV18 transcripts (Figure 2A). Of the 34 sites identified in the WT sample, 26 were also detected in the IVT sample (Figure 6A), suggesting that many of these calls may reflect background signal. When comparing stoichiometry between WT and IVT samples for the sites detected in WT, most sites clustered near the identity line, with only a few showing potentially higher stoichiometry in WT (Figure 6B, highlighted in red). Interestingly, three of these WT-enriched m^5^C calls were located exactly one nucleotide downstream of three of the strongest m^6^A sites; that is, at the C immediately following the DRACH A at positions 551, 598 and 618 (Figure 6C). This raised the possibility that nearby m^6^A may have confounded the m^5^C model, owing to altered current signal at the adjacent upstream base. Consistent with this interpretation, these three calls were not detected in the IVT data, where the adjacent m^6^A signal is also absent (Figure 6C, lower track).

**Figure 6.**
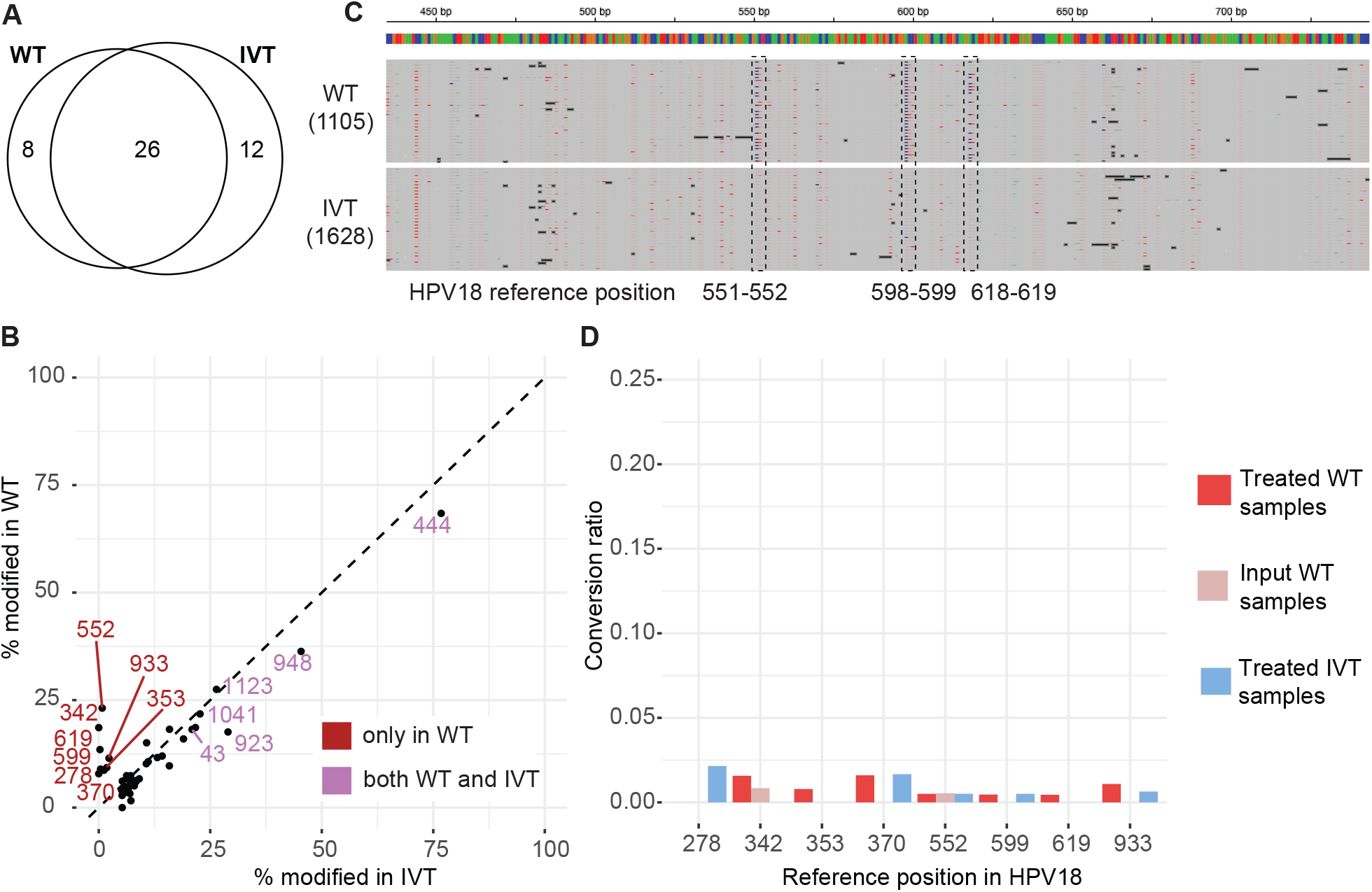
Minimal evidence for m^5^C modifications on HPV18 transcripts. (**A**) Venn diagram showing overlap of Dorado-identified m^5^C sites between the WT and IVT samples. (**B**) Scatter plot comparing m^5^C modification stoichiometry in WT vs. IVT at all sites called in WT. (**C**) IGV snapshot illustrating an example of a false-positive m^5^C call. In the WT reads, Dorado calls an m^5^C at cytosines (indicated by red marks) immediately following three of our identified m^6^A sites (blue marks). (**D**)m^5^C-TAC-seq results for the eight Dorado WT-specific m^5^C candidate sites. No positions showed statistically significant conversion ratios compared to control samples (based on a binomial generalised linear model).

To test whether HPV18 transcripts carry m^5^C, we reanalysed an orthogonal sequencing dataset, m^5^C-TAC-seq, from HeLa cells (Lu et al., 2024). In m^5^C-TAC-seq, methylated cytosines are detected through TET-assisted oxidation and chemical labelling, and are ultimately read out as C-to-T transitions in sequencing data. We reprocessed the HeLa WT and IVT control samples and examined cytosine positions on HPV18 for evidence of elevated conversion ratios relative to those observed in the input and IVT control samples. Special attention was given to the eight m^5^C sites detected by Dorado as WT-specific in the direct RNA sequencing data, all of which had adequate coverage in the m^5^C-TAC-seq samples (Figure S8A). This analysis showed only weak conversion ratios in the treated WT samples, which in all cases were not significantly different from the controls (Figure 6D). Thus, the chemical-labelling assay did not support m^5^C at any HPV18 site, supporting the interpretation that the Dorado m^5^C calls were false positives under these conditions.

Finally, for pseudouridine (Ψ), of the 26 sites detected by Dorado with at least 25 reads of coverage and 5% stoichiometry, 23 were also detected in the IVT sample (Figure S8B). Comparing stoichiometries between WT and IVT, all points lay close to the identity line, leaving no clear candidates for further analysis (Figure S8C) and providing little evidence for Ψ in the covered HPV18 transcript regions under the analysed conditions. Although we cannot rule out the presence of these modifications in regions with very low direct RNA sequencing coverage, our analyses overall provide little evidence for m^5^C or Ψ sites in HeLa-expressed HPV18 transcripts.

## Discussion

Nanopore long-read sequencing is increasingly being applied to study HPV-associated cancer biology. For example, targeted nanopore approaches have been used to map viral-host integration events in cervical tumours (Parida et al., 2024). Here, we used direct RNA nanopore sequencing with IVT controls and orthogonal chemical-profiling datasets to define a candidate map of RNA modifications on HeLa-expressed HPV18 early transcripts. Because DRS modification calling remains sensitive to model assumptions, sequence context, stoichiometry thresholds and local coverage, our combined approach allowed us to prioritise consistent candidates while reducing likely method-specific calls. The strongest evidence was obtained for m^6^A, where a set of sites was detected in native DRS but was largely absent from IVT controls, and was further supported by GLORI and eTAM-seq datasets. By contrast, we found little evidence for m^5^C or Ψ in the HPV18 regions that were sufficiently covered in the available datasets.

Among the identified m^6^A sites, position 598 in the E7 region emerged as the strongest candidate. This site showed high methylation in DRS, was supported by both GLORI and eTAM-seq, and decreased after STM2457 treatment, WTAP knockdown and FTO overexpression, supporting it as a well-supported m^6^A candidate deposited by the canonical methyltransferase complex. This observation is notable in light of a recent report demonstrating that a strong m^6^A site on E7-containing viral transcripts increases RNA stability via the reader IGF2BP1 (**?**). Although the precise correspondence between this site and the previously reported HPV16 site remains to be determined, our data nominate HPV18 position 598 as an obvious target for future functional analysis of transcript stability and oncogene expression.

One of the most interesting findings from our analysis is the association between m^6^A modification and processing of the HPV18 E6 intron. Across long DRS reads, eTAM-seq and staged 4sU-GLORI, position 224, which lies in a GAACT motif immediately upstream of the splice donor, showed consistently higher methylation in intron-retained reads than in the spliced E6*I isoform. Because retention of the E6-coding intron favours production of full-length E6 mRNA, whereas splicing generates the E6*I/E7 transcript required for efficient E7 expression, this places position 224 within an RNA-processing context that is likely to be relevant for high-risk HPV early gene regulation. In addition, because the unspliced transcript form was considerably rarer than the spliced form, this result illustrates how bulk measurements of m^6^A can mask important differences in rarer transcript sub-populations.

Recent HPV16 literature also supports the idea that m^6^A-related pathways can modulate early viral RNA processing. In particular, overexpression of the m^6^A-related factors ALKBH5 and YTHDC1 promoted retention of the E6-coding intron in HPV16, whereas ALKBH5 knockdown had the opposite effect, and METTL3 showed stronger effects on E1/E2-region splicing (Cui et al., 2022). In HPV18, our analyses of E6 intron retention and splicing indices in the eTAM-seq, 4sU-GLORI and GLORI perturbation datasets are broadly consistent with an association between m^6^A-related RNA processing states and E6 intron splicing, although variable coverage and reliance on short reads limits stronger mechanistic conclusions.

These HPV-specific observations are also consistent with broader principles linking m^6^A to RNA processing. YTHDC1 can regulate splice-site choice in selected transcripts (Xiao et al., 2016; Ke et al., 2017). In addition, recent work indicates that many m^6^A sites accumulate post-transcriptionally, become enriched in later nuclear RNA-processing states, and may promote nuclear retention of RNAs carrying intact 5^*′*^ splice-site motifs (Tang et al., 2024; Lee et al., 2025). By contrast, exon-junction-complex deposition can locally restrict METTL3-mediated m^6^A addition near exon–exon boundaries (Yang et al., 2022; Uzonyi et al., 2023; Liu et al., 2023). Within that framework, the higher methylation observed on intron-retained HPV18 molecules may reflect prolonged residence of unspliced or nuclear-retained RNAs, allowing greater access to the m^6^A writer complex, whereas successful splicing and exon-junction-complex deposition may subsequently limit further methylation near the junction. However, additional direct functional testing will be required to resolve the causal direction and underlying mechanisms.

In contrast to the E6 intron, we did not observe notable m^6^A differences associated with other viral RNA processing events. For example, inclusion of the E1 region or choice of the cryptic transcription initiation site previously observed by Yu et al. (2024) (+24, SV2) did not show differential m^6^A patterns in our data. However, our coverage of regions outside E6/E7 was limited, reducing our power to detect modifications there. Our re-analysis of GLORI and eTAM-seq data also suggested a small number of additional low-stoichiometry HPV18 m^6^A candidates that were not detected in the DRS data. These may merit future analysis, especially on late transcripts or in other sample conditions. We also note that nanopore reads often truncate at the 5^*′*^ end, which can limit confident distinction of transcripts arising from nearby initiation sites and reduce power to compare their modification levels.

A key strength of this study is that the same HPV18 m^6^A candidates recur across independent datasets generated in different laboratories and with different chemistries. At the same time, because the datasets reanalysed in this study were generated from independently propagated HeLa cells rather than from a single matched source, direct quantitative comparison of stoichiometries should be treated cautiously (Liu et al., 2019). In addition, the DRS analysis was based on one WT and one IVT sample, reflecting the constraints of available public data, and read coverage was limited for less abundant transcript classes, particularly the unspliced E6-containing molecules. Finally, HeLa cells harbour an integrated, rearranged HPV18 genome, so the findings may not fully generalise to infections with intact episomal HPV18 genomes (Yu et al., 2024). Despite these constraints, our study provides a useful starting point for understanding RNA modifications in HPV-driven cancers and generates focused, testable hypotheses for future work in matched experimental systems, additional models, and patient-derived material.

## Conclusion

Using independently generated public DRS, GLORI, eTAM-seq and 4sU-GLORI datasets, we generated a candidate map of m^6^A sites on HeLa-expressed HPV18 early transcripts, but find little evidence for m^5^C or pseudouridine in the covered HPV18 transcript regions under the conditions analysed. These results provide a focused resource for future matched experimental validation of viral RNA modifications and their potential roles in HPV18 gene regulation.

## Methods

All downstream analyses of processed data were conducted in R using custom scripts, unless otherwise stated. All plots, unless otherwise stated, were generated using the R package ggplot2. IGV browser was used to inspect reads and generate screenshots (Thorvaldsdóttir et al., 2013). P-values were adjusted using the Benjamini-Hochberg false discovery rate procedure, unless otherwise stated.

### Direct RNA sequencing analysis with Dorado

#### Dorado raw data processing

Direct RNA sequencing data for HeLa and associated IVT samples were downloaded from (Zou et al., 2025). All DRS data were basecalled with Dorado v0.9.0 using the most recent high-accuracy RNA model available at the time of analysis, rna004_130bps_sup@v5.1.0. During basecalling, reads were aligned to a reference consisting of HPV18 transcripts (Yu et al., 2024) merged with GRCh38 transcripts (GENCODE v47), using minimap2 with default settings as implemented in Dorado. Reads with a primary alignment to an HPV transcript and mapping quality *>* 5 were then remapped to the HPV18 genome using minimap2 v2.28 with options `-ax splice -uf -k14 -t 16 -y --secondary=no`.

#### Dorado modification calling

Basecalling was also performed with the available modification models `m$^{6}$A_DRACH, pseU and m5C`. Predicted modification probabilities were subsequently analysed with Modkit v0.4.2 using modkit pileup to tabulate position-level stoichiometries and `modkit extract full` to quantify modified sites per molecule. Only positions detected by Dorado with at least 5% stoichiometry and at least 25 mapped reads were retained as candidate sites for further analyses.

#### Read summaries

Read lengths, median lengths and coverage were computed excluding soft-clipped regions, using samtools, dorado summary and custom bash scripts.

#### Splice variants

Reads primarily aligned to HPV18 were assigned to six splice variants (SV1–SV6) using bedtools according to (i) 5^*′*^ end coordinate of the read, (ii) splicing of the E6 intron, identified by a 180–184N operation in the CIGAR string, and (iii) presence of E1 sequence. Read IDs were then used to create six separate BAM files for each of the WT and IVT samples. For the analysis of spliced versus unspliced reads, SV2–SV4 were merged as the spliced group and SV5–SV6 as the unspliced group. For the analysis of presence versus absence of E1, SV3 and SV6 were merged as E1-present and SV2, SV4 and SV5 as E1-absent. All BAM files were inspected in IGV before downstream analyses.

#### Analysis of spliced versus unspliced read sets

For each site, Fisher’s exact test was performed on a 2 × 2 table of counts (either spliced/unspliced or WT/IVT versus modified/unmodified). Only positions significantly more modified in WT than IVT (adjusted p-value < 0.01) were retained for subsequent tests of differential modification between spliced and unspliced read sets.

#### Sequence analysis

The R package GenomicRanges, together with BSgenome and the HPV18 genome reference, was used to extract 5-mer sequence context at candidate m^6^A sites (Lawrence et al., 2013; Pagès, 2018).

### Direct RNA sequencing analysis with xPore

#### Eventalign

All eventalign files were generated with the Nanopolish re-implementation f5c v1.5 to allow analysis of RNA004 data. Before applying `f5c eventalign`, raw data were converted from pod5 to the more efficient BLOW5 format using the python command `blue-crab p2s` followed by `slow5tools` merge, and were then indexed with `f5c index` (Samarakoon et al., 2023). Reads mapped to HPV18 (basecalled with Dorado as described above) were converted to FASTQ using samtools. Reference genome, BLOW5 files, FASTQ files and corresponding BAM files were passed to `f5c eventalign with options --signal-index, --rna, --scale-events, --min-mapq 0` and `--kmer-model rna004.nucleotide.5mer.model`.

#### xPore

Using the f5c-generated eventalign files, xpore dataprep was run before xPore v2.0 diffmod analysis. xPore was run with the RNA004 5-mer table provided through the xPore RNA004 branch and used as the prior in the config.yml file. Default xPore stopping criteria and maximum iterations were used (0.00001 and 500, respectively).

#### Downstream filtering

As xPore does not explicitly model genetic variation, all positions within ±4 nt of a SNP were removed from the xPore output. SNPs with *>*10% frequency were extracted from basecalled reads using samtools pileup. Positions inferred as modified were further filtered using a p-value threshold of p < 1e-7, as suggested previously (Tan et al., 2024b). P-value adjustment in Figure S2C was used only to break zero-defined p-values for plotting.

### GLORI data analyses

#### Data preprocessing

Raw sequencing reads for all HeLa GLORI samples (both GLORI+ samples involving the chemical protocol and GLORI-samples not treated by the chemical protocol) were downloaded from NCBI using SRA-tools (Leinonen et al., 2011). For GLORI+ samples, a combined GRCh38 + HPV18 reference was computationally converted from A to G and indexed using STAR v2.7.11a. Reads were mapped with STAR using options --clip3pAdapterSeq AGATCGGAAGAGC, --clip3pAfterAdapterNbases 0, --outFilterMismatchNoverReadLmax 0.10, --outFilterMismatchNmax 999, --outFilterMultimapNmax 50, --outFilterScoreMinOverLread 0, --outFilterMatchNminOverLread 0, --chimSegmentMin 15, --chimOutType WithinBAM SoftClip, --sjdbOverhang 74 and --outSAMattributes NH HI AS nM NM MD. Mapped reads were filtered for minimum mapping quality 20, filtered for alignment to HPV18, and samtools mpileup (Li et al., 2009) was used on all positions in the HPV18 reference genome.

#### Identification of HPV18 sites

For all adenosines in the HPV18 reference, counts of G (converted) and A (protected) for GLORI+ samples were used to calculate the protected-A proportion, A_count/(A_count + G_count). For GLORI-samples, the background A-to-G conversion ratio was calculated as G_count/(A_count + G_count). For assessing modified proportions at candidate sites in either GLORI+ or GLORI-samples, counts were first pooled across the two replicates of each control condition for the writer knockdown or inhibition experiments, the YTHDF knockdown experiments, or the hypoxia treatments (labelled siwritercontrol, siYTHDFcontrol and hypoxia-, respectively). Individual positions were retained if they had at least 5% protected-A ratio across pooled control samples, at least 5% protected-A ratio in both replicates of at least one control experiment, and negligible A-to-G mutation rates in the corresponding GLORI-samples. Variance at each position was calculated as np(1-p), where n is pooled read coverage at the position and p is the proportion of protected As.

#### Analysis of knockdown, inhibition or hypoxia experiments

Control samples for the m^6^A writer-complex perturbation experiments (siwritercontrol) were compared with each perturbation (siMETTL3, siMETTL14, siWTAP or STM2457+), YTHDF knockdown samples were paired with their controls (siYTHDFcontrol), and hypoxia-treated samples (hypoxia+) were paired with hypoxia controls (hypoxia-). To assess positions with significantly reduced protected-A ratios upon perturbation, a generalised linear model with binomial family and condition (control or perturbed) as covariate was fitted using both replicates for each condition. Adjusted p-values were converted to −log_10_ scale before plotting with the R package pheatmap.

#### Analysis of GLORI 4sU datasets

Staged nuclear 4sU-GLORI reads from HeLa cells (Tang et al., 2024) were processed using the same mapping and protected-A quantification framework described above for GLORI. Briefly, reads from the early 4sU+ poly(A)-, mid 4sU+ poly(A)+ and late 4sU-poly(A)+ nuclear RNA fractions were adapter-trimmed with cutadapt, mapped with STAR to the A-to-G-converted GRCh38+HPV18 reference, and filtered for primary HPV18 alignments with mapping quality ≥20 and aligned length ≥25 nt. Samtools mpileup was used to extract A and G counts at HPV18 adenosines, and protected-A fractions were calculated as A/(A+G).

### Analysis of eTAM-seq data

Raw eTAM-seq reads from HeLa FTO- and FTO+ samples were downloaded from GSE201064 and processed with HPV18 included in the reference genome. A HISAT-3N index was built on a combined GRCh38+HPV18 reference with `--base-change A,G` and splice/exon annotations from the corresponding combined GTF. FASTQ files were merged across lanes, adapter-trimmed in two rounds with cutadapt, and a 6-nt UMI was extracted from read 2 using UMI-tools. Reads were aligned to the combined reference with HISAT-3N using `--base-change A,G` and `--rna-strandness F`, filtered by mapping quality, deduplicated with UMI-tools, and filtered for sufficient A-to-G conversion. HPV18-mapped reads were then extracted and analysed with pileup2var (-a A) to obtain per-position A and G counts. Protected-A proportions were calculated at HPV18 adenosines as A/(A+G).

### Intron retention and linkage analyses

For analyses of E6 intron retention and read-level linkage at positions 224 and 455, reads covering the candidate position were classified as E6*I-spliced if they contained the E6*I junction in the CIGAR string within the accepted donor–acceptor window, classified as intron-retaining if they overlapped the intron without that junction, and otherwise excluded as uninformative. Protected-A fractions were then recalculated separately for the spliced and intron-retaining read classes. Where reported, intron retention indices were calculated as unspliced / informative reads.

### m^5^C-TAC-seq analysis

#### Data processing

Raw m^5^C-TAC-seq reads were downloaded using SRA Toolkit. Low-quality bases and adapter sequences were trimmed using cutadapt (Martin, 2011). R2 reads were deduplicated using UMI information (Smith et al., 2017), clipped by 10 nt using seqkit (Shen et al., 2016) and fastp (Chen et al., 2018), and mapped to the combined GRCh38 and HPV18 genomes using HISAT-3N (Zhang et al., 2021). C-to-T mismatches were extracted from mapped BAM files and summarised using the `hisat-3n-table` command.

#### Downstream analyses

A custom R script was used to process and analyse C-to-T conversion ratios at HPV18 cytosines. To test for evidence of m^5^C-specific conversion in treated WT RNA, we fitted a binomial generalised linear model using the formula `cbind(converted, unconverted) ∼ RNA_type * treatment, where RNA_type` denotes WT or IVT RNA and treatment denotes input or chemically treated samples. The interaction term was used to assess whether chemically treated WT samples showed elevated conversion relative to the corresponding controls.

## Data Availability

Nanopore direct RNA sequencing data for HeLa cells were downloaded from the ENA under accession PRJEB80229. Raw GLORI sequencing data were downloaded from the NCBI Gene Expression Omnibus (GEO; GSE210563), raw eTAM-seq reads from GEO accession GSE201064, and raw m^5^C-TAC-seq reads from GEO accession GSE242724. Raw 4sU-GLORI reads were downloaded from the Genome Sequence Archive for Human database under accession HRA006964, associated with project PRJCA024492. The reference sequence for the HPV18 genome was downloaded from NCBI (Reference Sequence: NC_001357.1).

## Author contributions

I.R.H analysed the DRS and GLORI HeLa data with support from S.R, S.R analysed the eTAM-seq data, M.S.L.S analysed the GLORI 4sU data, S.R. supervised the project with input from J.J and J.Y.A. S.R. wrote the manuscript with input from all authors.

## Supplementary Figures

**Figure S1.**
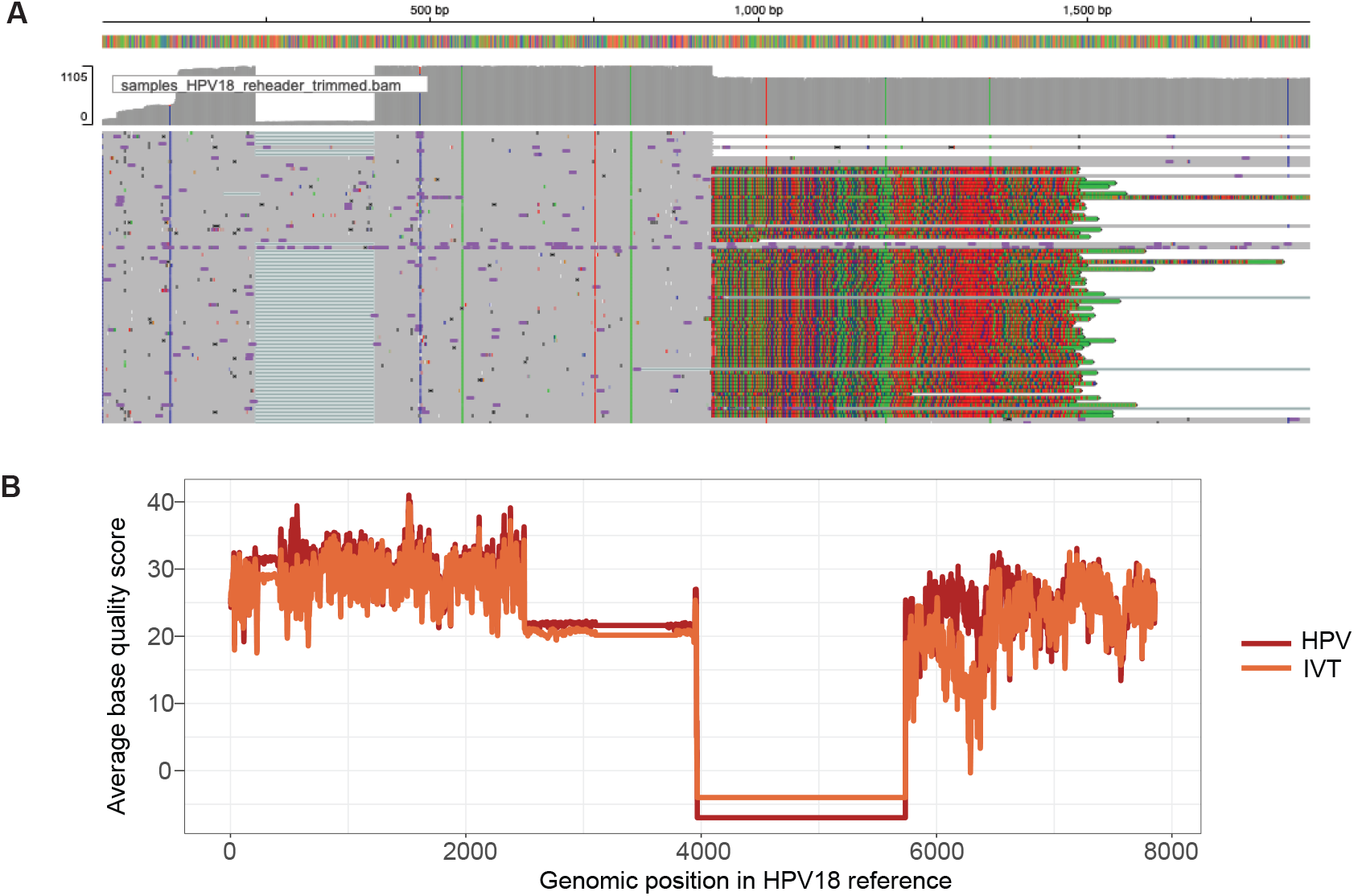
Quality check of HPV18-mapped reads using direct RNA-sequencing. (**A**) IGV screenshot illustrating the presence of softclipped (multi-coloured) regions in reads, derived from the human genome. (**B**) Average base quality per position for each of the HPV and IVT samples across the HPV18 reference genome. Zero quality indicates lack of coverage in these regions.

**Figure S2.**
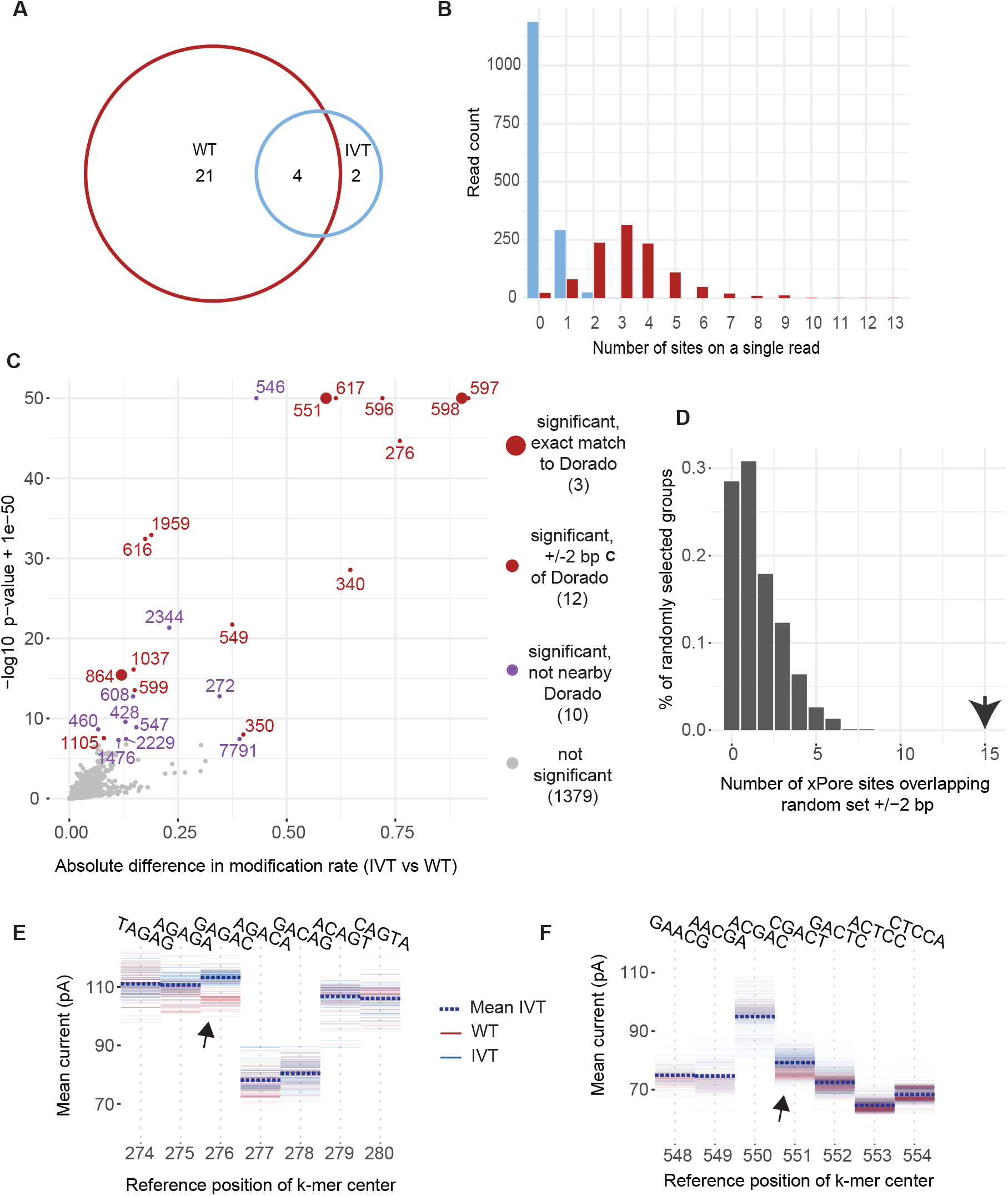
Properties of m^6^A sites identified by Dorado in the WT and IVT samples. (**A**) Venn diagram showing overlap between WT and IVT samples for Dorado-identified sites. (**B**) Distribution of sites per molecule for the 25 WT-derived sites, shown for the WT and IVT samples. (**C**) xPore results comparing WT and IVT samples. Each point represents a genomic position, plotted by xPore’s inferred modification-rate difference (WT–IVT, absolute value) versus *−*log_10_(adjusted p-value). Points are coloured by xPore significance (purple for FDR < 0.05, grey for not significant). Red circles denote positions corresponding to Dorado m^6^A candidates (large red circles for exact nucleotide matches; small red circles for positions within *±*2 nt of a Dorado-called site). (**D**) Distribution of xPore sites overlapping sets of 25 randomly selected positions. The arrow represents the true overlap between the xPore significant sites and the Dorado-identified sites (*±*2 bp). (**E-F**) f5c eventalign signal plots for WT (red) and IVT (blue) reads around two Dorado-identified m^6^A candidate sites, with the xPore-identified positions indicated by black arrows: (**E**) position 277 and (**F**) position 551, including 3 bp upstream and downstream of the site of interest. Thick dotted horizontal lines show the mean signal level for the IVT sample, providing a reference reflecting the IVT baseline.

**Figure S3.**
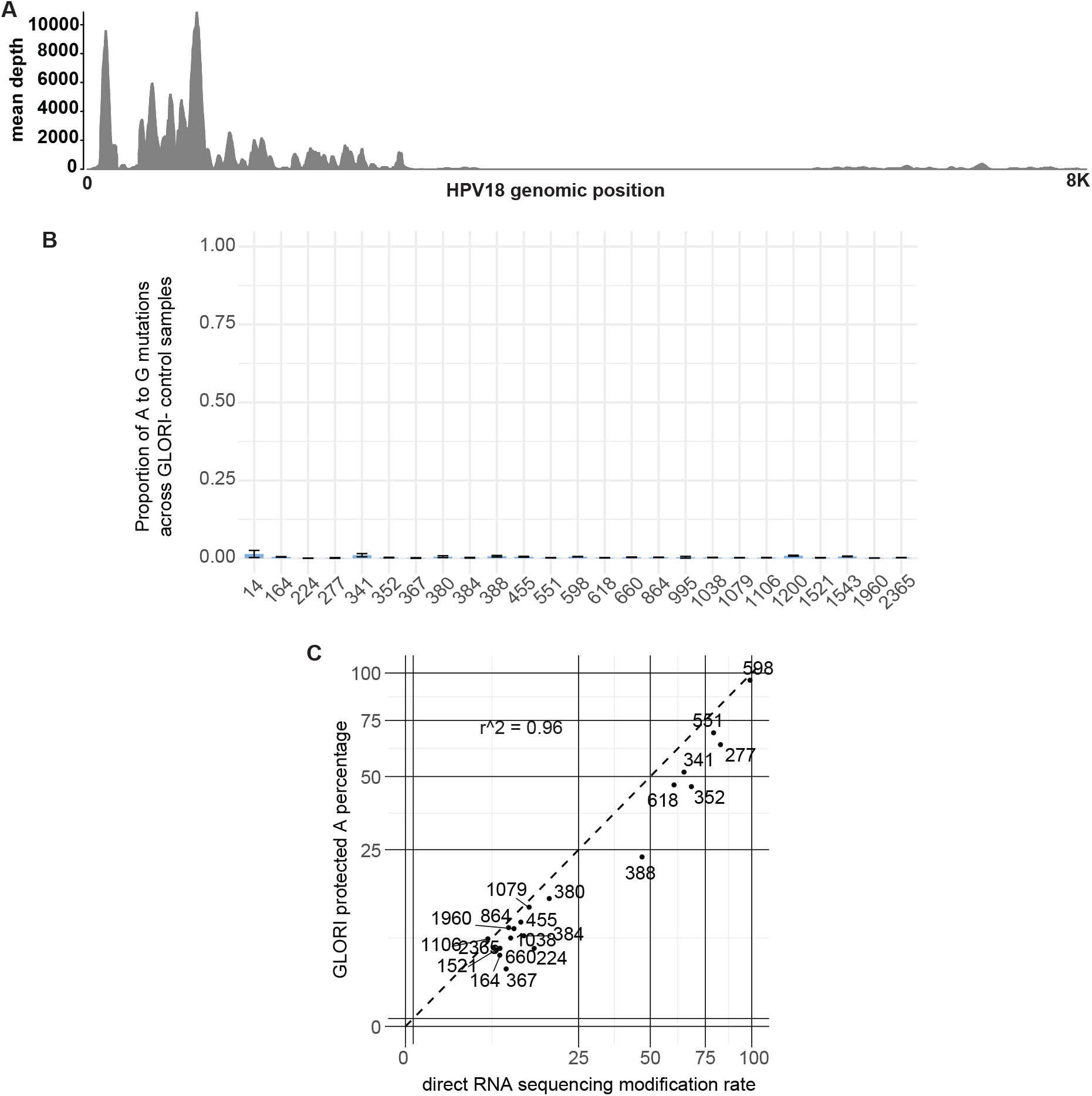
GLORI quality control analyses. (**A**) Coverage of GLORI reads across the HPV18 genome, based on wild-type control samples for the knockdown, inhibition and hypoxia experiments. (**B**) Proportions of A-to-G conversions in untreated GLORI (GLORI-) samples for the knockdown, inhibition and hypoxia experiments, for each of the 25 candidate m^6^A sites identified by Dorado. (**C**) Scatter plot comparing estimated modification rates from nanopore direct RNA sequencing and GLORI for the 21 m^6^A sites identified only in the WT sample.

**Figure S4.**
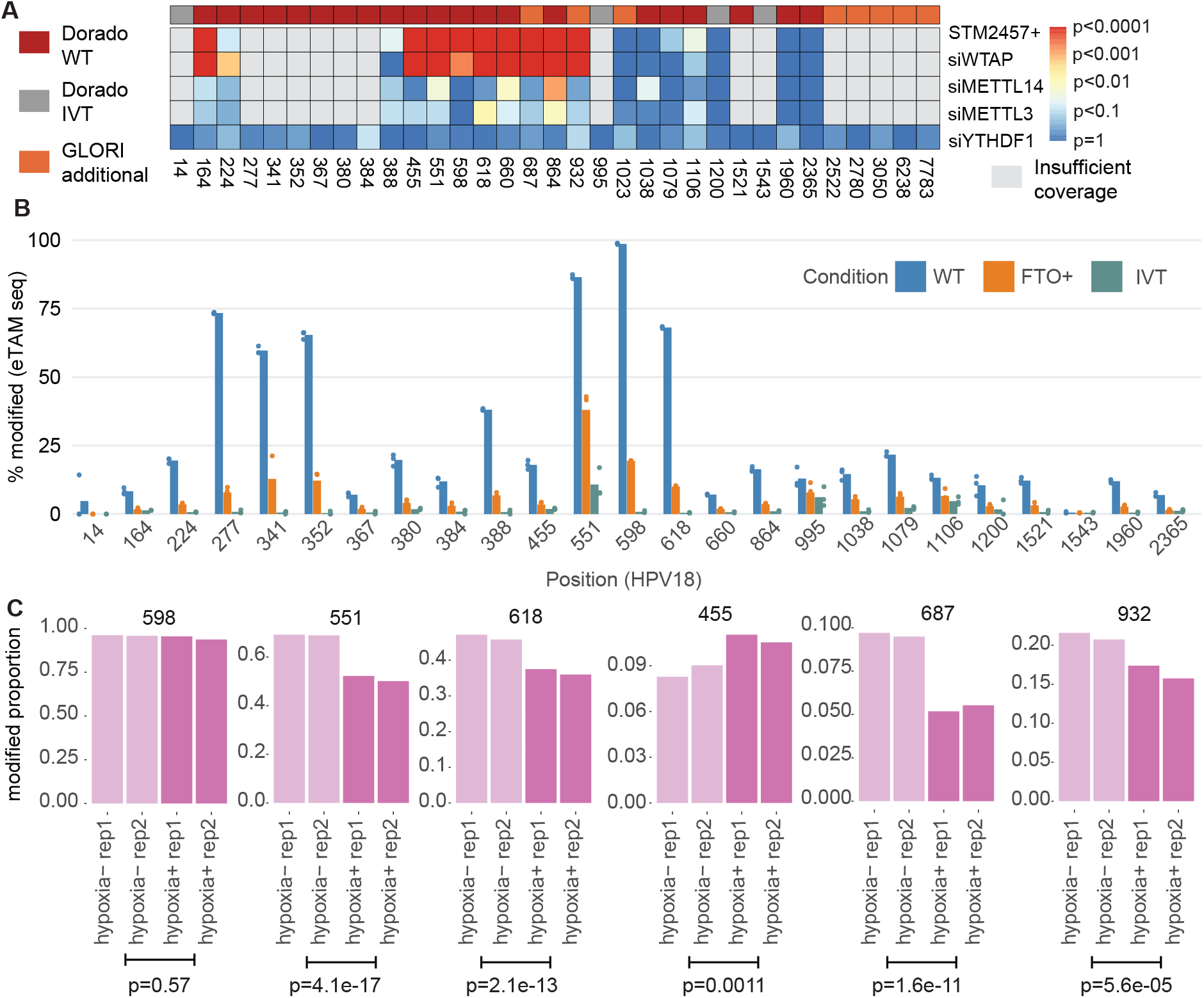
GLORI analysis of methylation changes based on knockdown, inhibition or hypoxia experiments. (**A**) Heatmap showing significance of writer component knockdown or inhibition versus controls (siwriterControl) or of YTHDF2 knockdown versus controls (siYTHDF2control). Adjusted p-values from the binomial generalised linear model were converted into -log10(.) prior to plotting the heatmap. Coloured bar at the top represents the three groups indicated to the left of the plot. (**B**) Extended eTAM figure for DRS-defined sites, including the adenosine ratios detected in the IVT samples. (**C**) Analysis of hypoxia samples. Bars represent proportion of protected As from treated GLORI samples, and p-values represent significance of hypoxia treatment based on a binomial generalised linear model.

**Figure S5.**
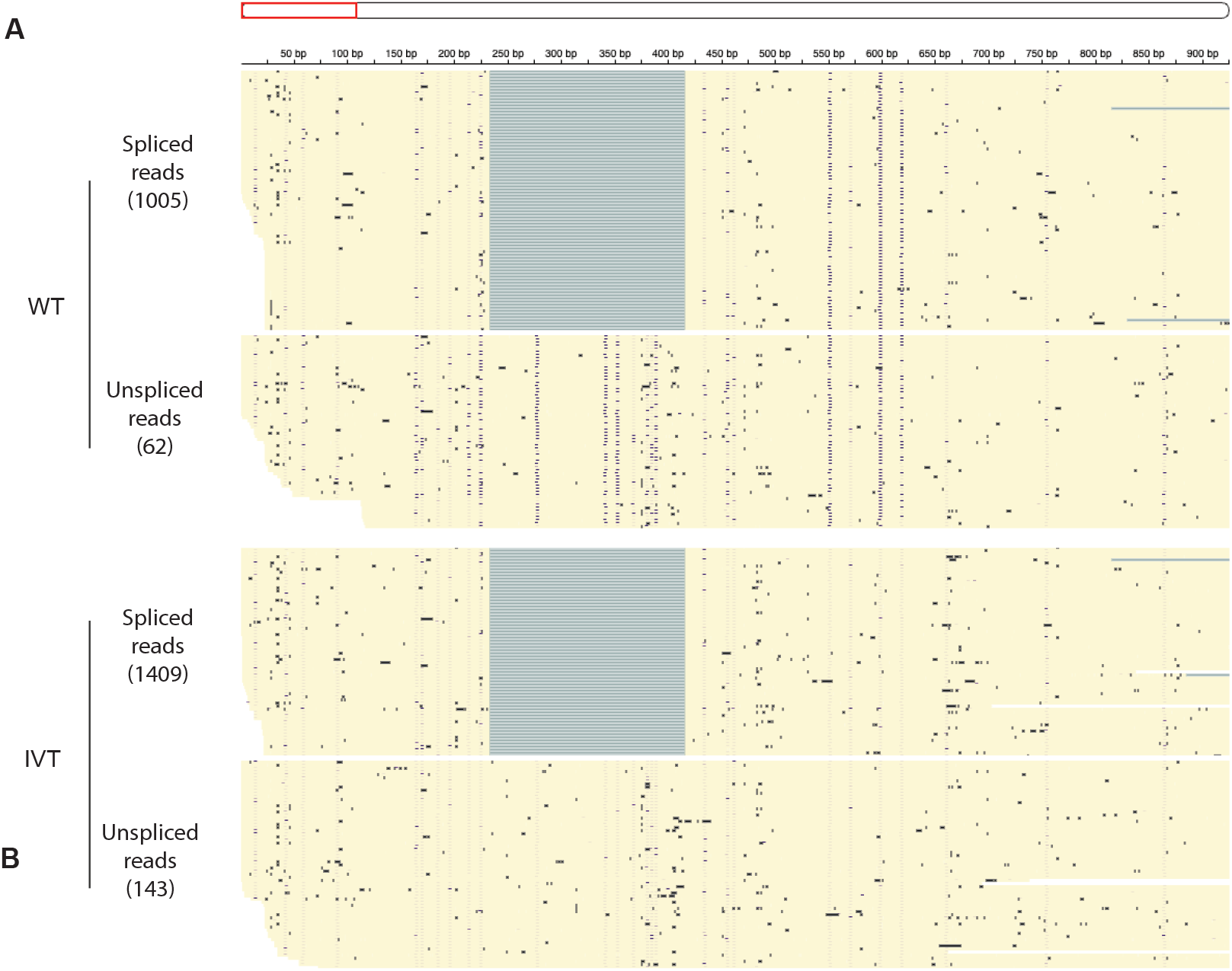
Extended IGV view of E6-intron spliced and unspliced reads. IGV view including spliced and unspliced reads from the WT sample. Blue dots represent Dorado m^6^A modification probabilities. The number of reads considered in each class is indicated on the left. (**B**) As in (**A**), but for the IVT sample.

**Figure S6.**
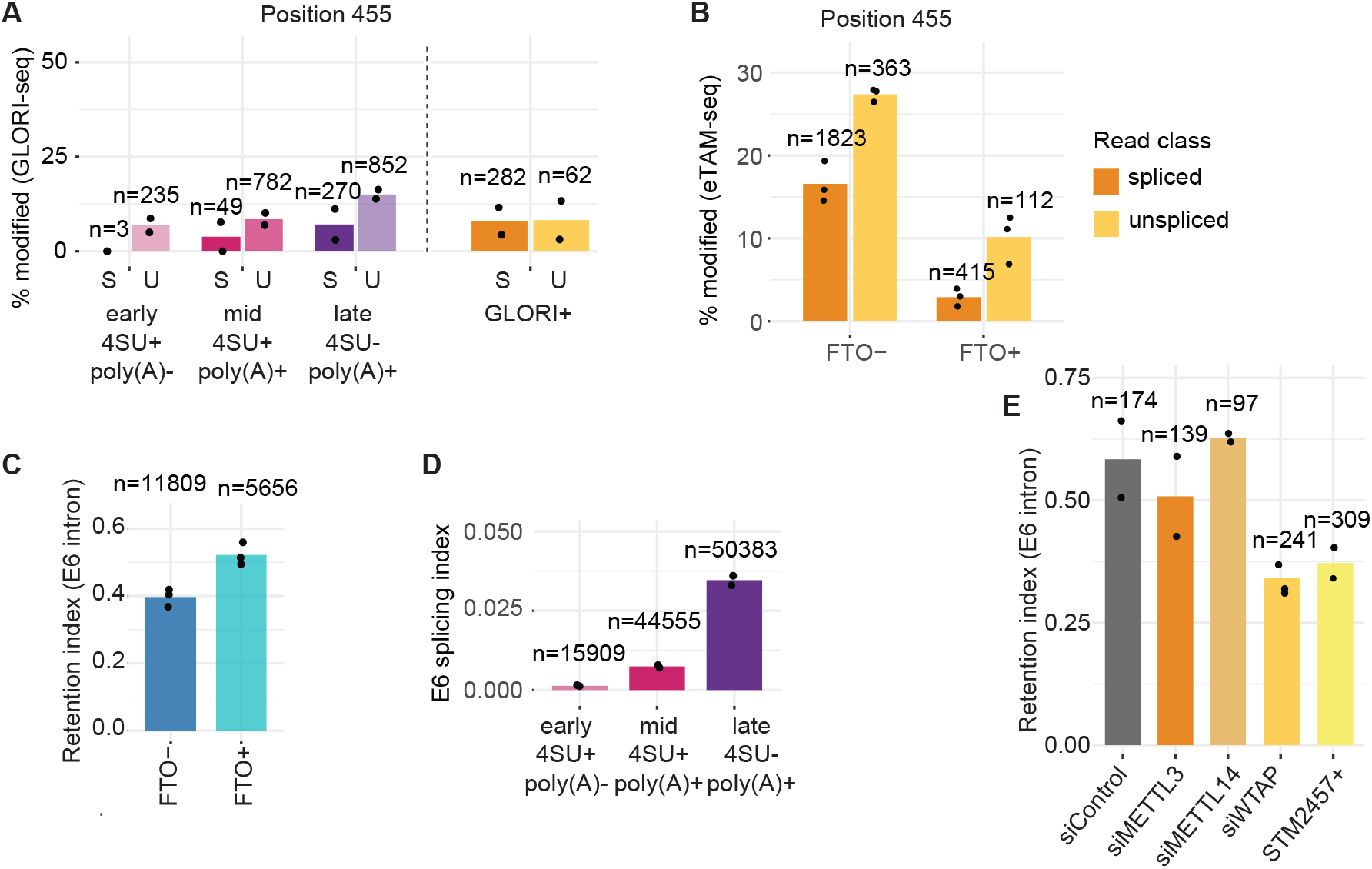
Additional analyses of E6 intron splicing based on eTAM-seq and GLORI datasets. (**A**) E6 intron retention (unspliced / informative reads) in eTAM-seq data. Dots represent values for individual replicates, and n values indicate numbers of informative reads. (**B**) Splicing index of the E6 intron (spliced / informative reads) in the staged 4sU-GLORI dataset. (**C**) As in (**A**), but for the GLORI writer-perturbation dataset. (**D**) Linkage analysis for the m^6^A site at position 455 in the staged 4sU-GLORI dataset and the GLORI HeLa hypoxia control sample. (**E**) As in (**D**), but based on eTAM-seq data.

**Figure S7.**
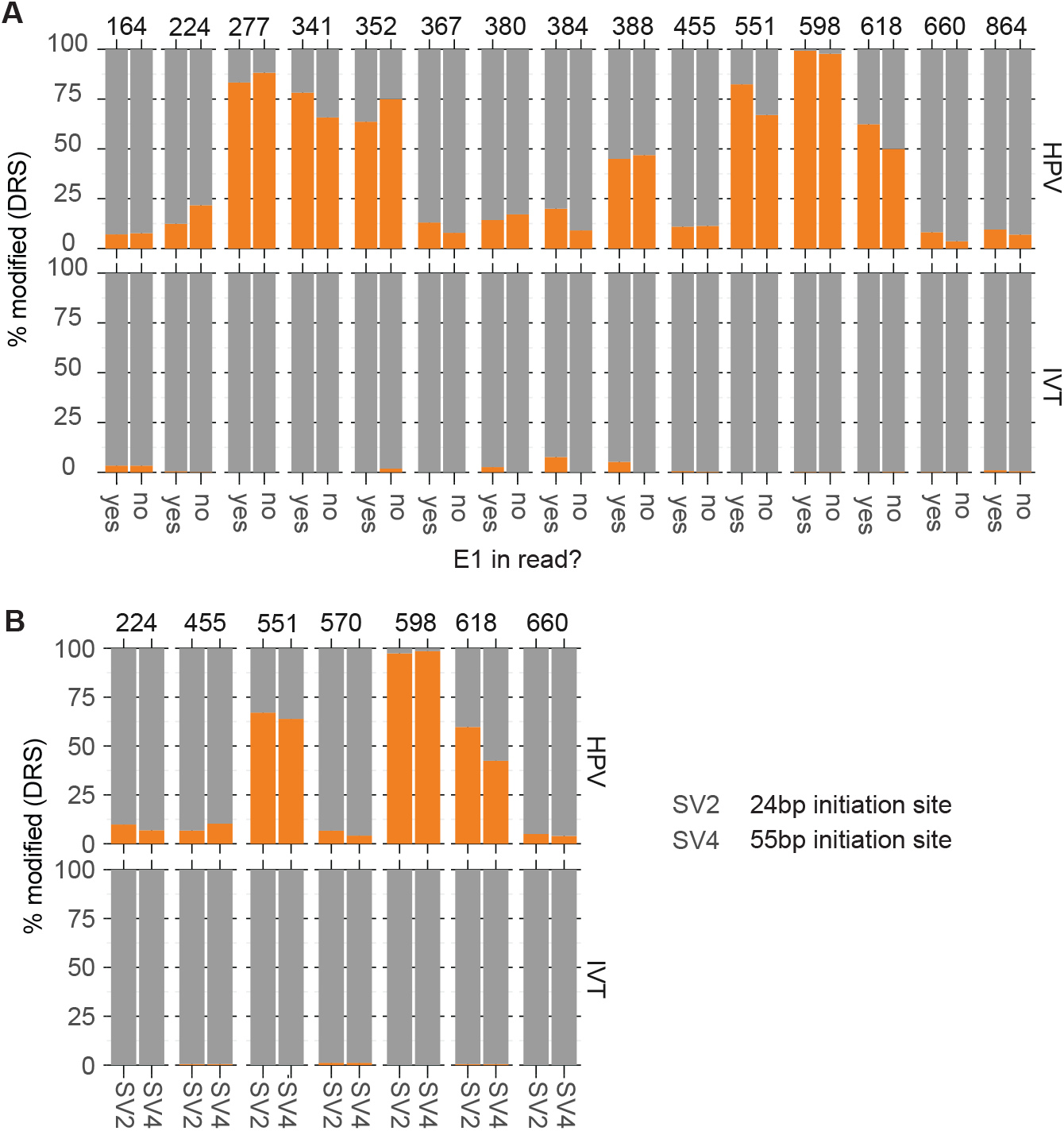
Analysis of differential methylation according to presence of E1 or use of transcription initiation site. (**A**) WT and IVT reads from direct RNA-sequencing split according to E1 presence, plotted is the modification rate inferred by modkit. (**B**) as (**A**) but reads for SV2 and SV4 are compared. All positions were tested using a Fisher’s exact test (based on counts yes/no versus modified/unmodified reads) and none were significant after multiple testing adjustments.

**Figure S8.**
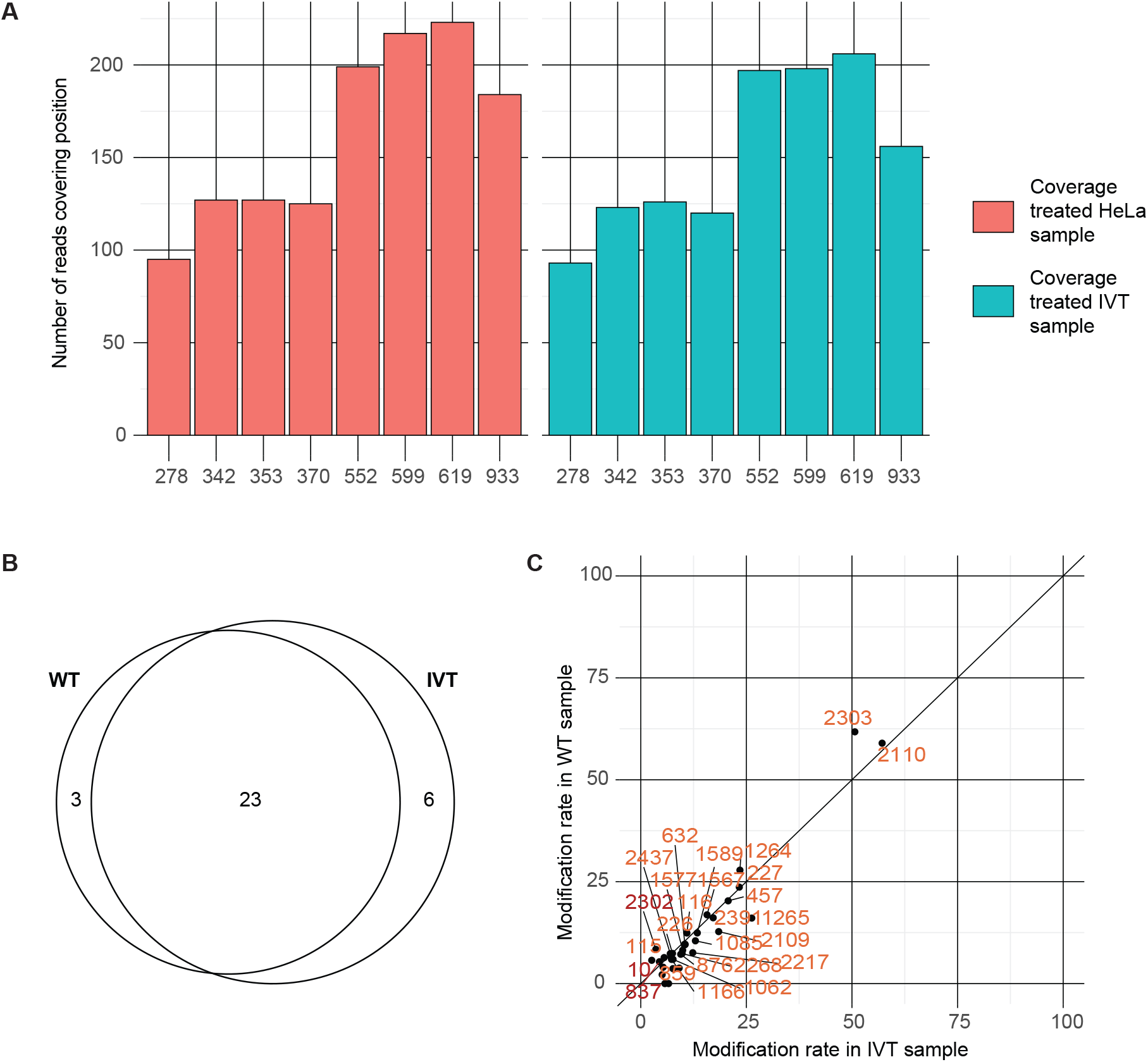
Analysis of m5C and Pseudouridine. (**A**) Coverage of m^5^C-TAC-seq reads in chemically treated samples at the 8 candidate positions which were detected in the WT direct RNA-sequencing sample but not the IVT (**B**) Overlap between WT and IVT Psi sites identified by Dorado (**C**) Stoichiometry scatter plot for Psi.

